# LoLoPicker: Detecting Low Allelic-Fraction Variants in Low-Quality Cancer Samples

**DOI:** 10.1101/043612

**Authors:** Jian Carrot-Zhang, Jacek Majewski

**Affiliations:** Department of Human Genetics, McGill University, Montreal, Quebec, Canada; McGill University and Génome Québec Innovation Centre, Montreal, Quebec, Canada.

**Keywords:** somatic variant calling, low allelic-fraction variants, FFPE tissues, high specificity

## Abstract

**Summary:** We developed an efficient tool dedicated to call somatic variants from next generation sequencing (NGS) data with the help of a user-defined control panel of non-cancer samples. Compared with other methods, we showed superior performance of LoLoPicker with significantly improved specificity. The algorithm of LoLoPicker is particularly useful for calling low allelic-fraction variants from low-quality cancer samples such as formalin-fixed and paraffin-embedded (FFPE) samples.

**Implementation and Availability:** The main scripts are implemented in Python 2.7.8 and the package is released at https://github.com/jcarrotzhang/LoLoPicker.

## Introduction

The detection of somatic mutations in tumors remains challenging. One of the major complexities is that variants with low allelic-fraction are commonly observed in tumor samples, and the difficulty of identifying those variants is magnified by the fact that sequencing technologies are still imperfect and error-prone. Moreover, technical artifacts, such as C to T and G to A transitions can arise from the formalin-fixation process and additional filters against FFPE-specific errors are required (Williams *et al.,* 1999, Yost *et al.,* 2012, Spencer et al., 2013).

NGS has emerged as a promising tool to discover disease-causing genes. For many basic research or clinical laboratories, the number of samples being sequenced has increased dramatically. Some laboratories build their in-house database to enable them filtering out false-positive calls that are specific to library preparation, protocols, instruments, environmental factors or analytical pipeline. MuTect also recommends using a panel of normal samples to filter missed germline variants and recurrent artifacts (Cibulskis et al. 2013). Moreover, a control panel provides an opportunity to precisely estimate the site-specific error rates that give the advantage to increase the sensitivity of calling single nucleotide variants (SNVs) on sites with lower error rates, and reduce false positives on sites with high error rates. This idea has been implemented for targeted re-sequencing experiments (Gerstung *et al.,* 2013).

Here we present LoLoPicker, a flexible variant caller that can handle low-quality samples. This program allows users to provide a control panel consisting of normal samples processed using similar procedures as the tumor (e.g. FFPE samples), and uses this control panel to identify artifacts or germ-line variants using a *K*-means clustering. Then, a binominal test is performed to determinate whether the ratio of reads supporting the tumor variant exceeds the background error rate (Figure 1). Detailed description of the algorithm is provided in the Supplementary Information file.

**Figure 1:**
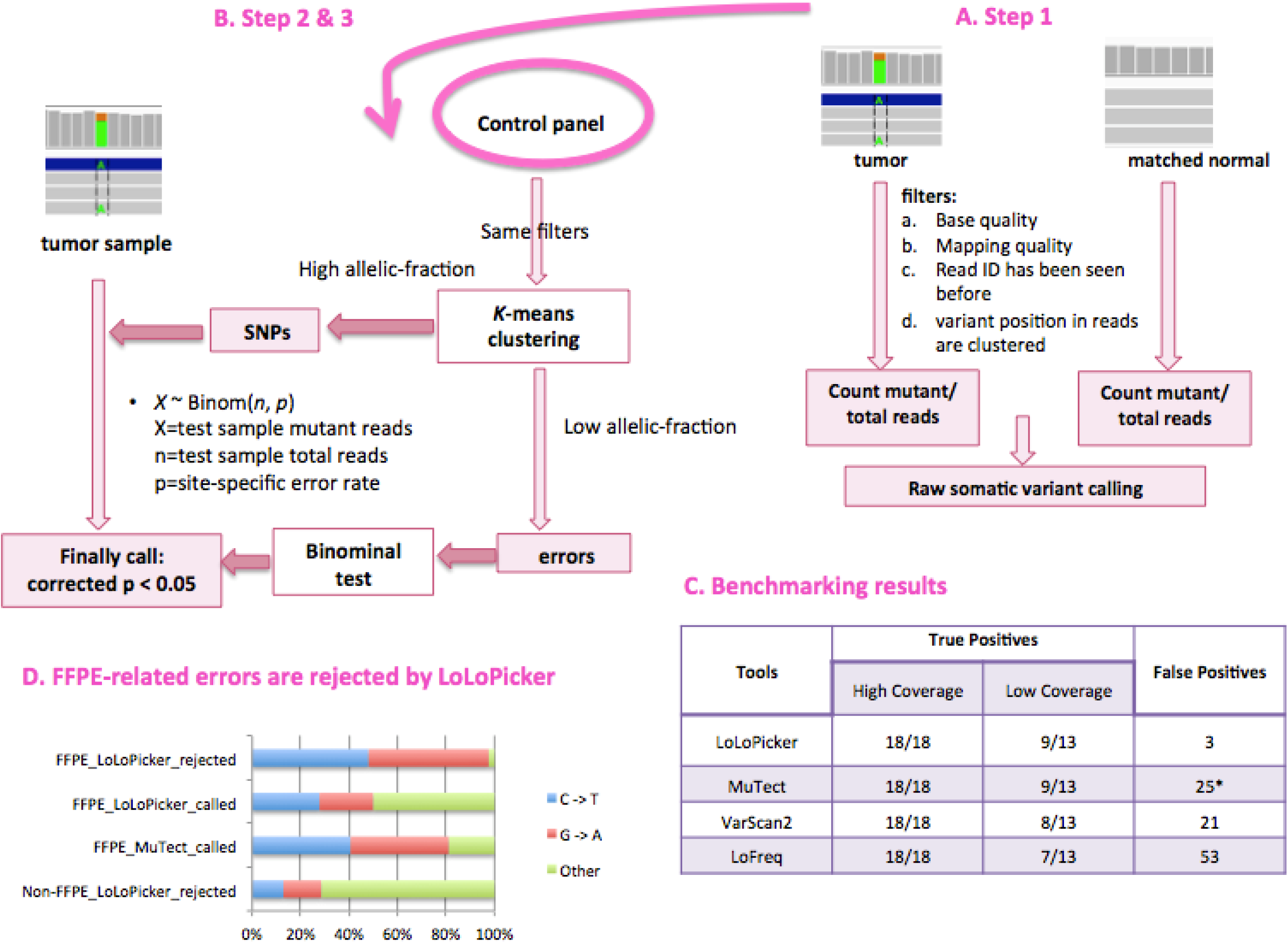
Workflow and performance of LoLoPicker. A. Step 1: LoLoPicker performs raw variant calling using tumor and matched normal sample. B. Step 2 and 3: LoLoPicker performs its core statistical framework using a user-provided control cohort. C. Number of true positives and false positives called by LoLoPicker, MuTect, VarScan and LoFreq from benchmarked samples, suggesting high sensitivity and specificity of LoLoPicker. D. C to T and G to A transitions, which are known errors enriched in FFPE samples compared to fresh-frozen samples, are frequently observed in LoLoPicker-rejected variants, and MuTect-called variants; whereas these transitions are less frequent among variants called by LoLoPicker. *False positives of MuTect are reduced to 15, when we apply the “‐‐normal_panel” option in MuTect using the same controls as LoLoPicker.

## Benchmarking Analysis

To access the performance of LoLoPicker in comparison to other variant callers, we benchmarked LoLoPicker, MuTect, VarScan2 and LoFreq against two datasets (Cibulskis *et al.,* 2013, Koboldt *et al.,* 2012, Wilm *et al.,* 2012). Somatic SNVs identified from whole-exome sequencing (WES) of an ovarian tumor and validated by Sanger sequencing were used as a set of true positives (Zhang *et al.,* 2013). BAM files of the tumor and its matched blood sample were mixed and down-sampled to ensure that variants are present in low allelic-fraction. For specificity, a sample that underwent WES twice was used, and all variants called between the two batches were considered as false positives. Moreover, 500 unrelated germ-line samples from non-cancer patients were used in our control panel. As the results, LoLoPicker showed highest sensitivity and specificity (Figure 1).

## Applying LoLoPicker to Real Data

### Fresh-frozen tumor samples

We then applied LoLoPicker, MuTect and VarScan2 to analyze a real cancer sample from a glioblastoma (GBM) patient (GBM_9). Known GBM driving mutations were identified, including mutations in *TP53*, *H3F3A*, *ATRX*, and *PIK3CA*. Only LoLoPicker successfully identified all of them. MuTect discarded the *TP53* mutation because it found three reads supporting the variant in the matched germ-line sample. In LoLoPicker, the mutation was retained because we count overlapping read-pair covering same variant as one (Figure S3). VarScan2 missed the *PIK3CA* mutation. Of note, the *PIK3CA* mutation showed low allelic-fraction at 6%. Again, this demonstrates that LoLoPicker has a high sensitivity of calling low allelic-fraction variants. Moreover, 14 SNVs in GBM_9 were selected for targeted re-sequencing validation. We found that variants called by both LoLoPicker and MuTect were all validated, whereas all variants that called by MuTect but rejected by LoLoPicker were not validated. These included four variants supported by higher coverage (>=5X) (Table S3). Our results suggested the high specificity of LoLoPicker without rejecting true positives as trade-off.

### FFPE tumor samples

Site-specific error rates in FFPE samples are much higher than fresh-frozen samples (Figure S6). In our previously published work on small cell carcinoma of the ovary, hypercalcemic type (SCCOHT), we showed that only one gene – *SMARCA4* – was recurrently mutated in the tumors, and no other mutations were responsible for this disease (Witkowski *et al*., 2014). We, therefore, tested LoLoPicker on an FFPE-SCCOHT sample (UN5). Although very few somatic mutations other than *SMARCA4* mutations were expected, both MuTect and VarScan2 called a large number of SNVs (483 and 143, respectively). Even though a normal panel of 35 FFPE samples was provided to MuTect, only 19 variants were filtered out by the “‐‐normal_panel” option. By contrast, using the same normal panel, LoLoPicker only called 18 variants. Most of the LoLoPicker rejected calls were C to T and G to A transitions known to be induced by the FFPE protocol (Yost *et al*., 2012, Spencer *et al*., 2013), suggesting the necessity of providing a control cohort to further reduce false positive calls related to batch effects, especially FFPE-specific artifacts (Figure 1).

## Discussions

LoLoPicker is tailored for low allelic-fraction, somatic SNVs. Compared to other programs, LoLoPicker showed highest sensitivity and more importantly, dramatically improved specificity, thus highlighting the importance of precisely measuring site-specific error rate from additional control samples, rather than from the matched normal sample solely. Although we expect that LoLoPicker will handle data from any sequencing platforms and alignment methods, we suggest that samples processed in similar experimental protocols and analytical pipelines should be used. For example, having a control panel of FFPE samples helped in reducing FFPE-specific artifacts. Compared to simply filtering out recurrent calls from normal samples, LoLoPicker’s statistical framework retains sites with lowlevel artifacts, allowing high sensitivity. Finally, the LoLoPicker algorithm can be easily parallelized to allow WES analysis against a lager number of control samples in a reasonable time, although we showed that 35 FFPE controls were able to reject most of the false positives. As FFPE is commonly used in clinical laboratories, our method will pave the way for NGS application in cancer research.

## Acknowledgements

We thank Hamid Nikbakht and Simon Papillon for their helpful discussions. Pierre Lepage for his help with targeted re-sequencing. JM is the recipient of a Canada Research Chair in Genomics.

